# PLK4 homodimerization is required for CEP152 centrosome localization and spindle organization

**DOI:** 10.1101/2023.11.06.565834

**Authors:** Harshita Kasera, Priyanka Singh

**Affiliations:** Department of Bioscience & Bioengineering, Indian Institute of Technology Jodhpur NH 62, Nagaur Road, Karwar 342037, Jodhpur, Rajasthan, India

**Keywords:** Centrosome, Centriole, PLK4, CEP152, CEP192

## Abstract

Polo-Like Kinase 4 (PLK4) is a unique serine/threonine kinase family member that homodimerizes using its cryptic polo-box (CPB) region. Homodimerization of PLK4 causes transphosphorylation, which activates its ubiquitin-mediated degradation. Interestingly, CPB is also involved in interaction with the upstream centrosome recruiters, CEP152 and CEP192 in human cells. However, the effect of PLK4 homodimerization on the CEP192-CEP152 network remains unexplored. In this work, we identified a frequently occurring cancerous PLK4 variant (E774*), which truncated the protein at the 774 position. The truncated PLK4 is unable to homodimerize and interact with CEP152 and CEP192. During the S-phase progression, we show that CEP152 recruits PLK4 to centrosomes. The homodimerization of PLK4, in turn, is needed for maintaining CEP152 centrosome levels. CEP152 levels correlate to pericentrin at S-phase centrosomes, which generate focused spindles by the M-phase. The homodimerization mutant exhibits reduced levels of CEP152 and pericentrin at S-phase centrosomes, which causes unfocused spindles at the M-phase and reduces cell viability. This work shows the requirement of PLK4 homodimerization for proper centrosome and spindle organization, which is disrupted in cancer.

## INTRODUCTION

Polo-Like Kinase 4 (PLK4), belonging to the serine/threonine polo-like kinases family (PLKs) (Barr *et al*, 2004), is localized to membrane-less organelles called centrosomes in dividing cells. A centrosome comprises a pair of cylindrical microtubule-based structures known as centrioles, surrounded by a protein matrix called pericentriolar material (PCM). Centrosomes play a significant role in cell division by forming spindle microtubules (Jaiswal *et al*, 2021). Any abnormality in the centrosome organization and number in a dividing cell could cause spindle defects, which leads to abnormal chromosome segregation and cancer (Godinho & Pellman, 2014; Ganem *et al*, 2009). Therefore, centrosome duplication is tightly regulated by the cues in the S-phase of the cell cycle.

PLK4 is a master regulator for centriole duplication events and dictates centrosome numbers (Habedanck *et al*, 2005). At the early G_1_-phase of the cell cycle, the centriole pair disengages but remains tethered *via* a proteinaceous linker. This relaxation of the two orthogonally juxtaposed centriole pairs licenses them for duplication in the S-phase of the cell cycle (Fujita *et al*, 2015). At this stage, PLK4 marks the location adjacent to the parent centriole, where the new centriole (procentriole) is generated (Fu *et al*, 2015). The overexpression of PLK4 induces rosettes of procentrioles around the parent centriole (Kleylein-Sohn *et al*, 2007). Accordingly, dysregulated levels of PLK4 have been associated with the onset and progression of many cancers (Liu *et al*, 2012; Shinmura *et al*, 2014; Wang *et al*, 2019; Marina & Saavedra, 2014).

The functional orthologue of PLK4 is known as ZYG-1 (Pelletier *et al*, 2006) in *Caenorhabditis elegans* and SAK/PLK4 (Habedanck *et al*, 2005; Bettencourt-Dias *et al*, 2005) in *Drosophila melanogaster*. It contains a conserved kinase domain at the N-terminal region and characteristic polo-boxes (PBs) at the C-terminal region (**Fig 1A**). The polo-box domain (PBD) of PLK4 shows a structural divergence from the originally identified and widely studied family member, the mitotic kinase PLK1. The PLK1 PBD, containing two PBs separated by a linker region, is involved in intramolecular dimerization, influencing its kinase activity. On the other hand, PLK4 PBD has three PBs, where the PB1 and PB2 regions are in tandem with each other, and together, they are referred to as the cryptic polo-box (CPB). This region is followed by a short linker and a third polo-box region (PB3). PLK4 CPB region is involved in centriole localization and intermolecular homodimerization. A 2.3 Å resolution crystal structure of *Drosophila* PLK4 CPB identified homodimerization with side-to-side winged architecture. This region is required for PLK4 centriole targeting *via* interaction with the Asterless (Asl, orthologue of human CEP152) protein (Slevin *et al*, 2012). Later, analysis of the X-ray crystal structure of the CPB domain of ZYG-1, the *C. elegans* functional orthologue of PLK4, at 2.54 Å resolution revealed an X-shaped end-to-end conformation (Shimanovskaya & Dong, 2014; Shimanovskaya *et al*, 2014). The dimerization of PLK4 triggers trans-phosphorylation of its phosphodegron motifs in the linker-1 region, which results in β-TrCP-mediated proteasomal protein degradation (Klebba *et al*, 2013, 2015; Cunha-Ferreira *et al*, 2013) and thus dimerization maintains a tight control over centriole number. However, recent work using *Drosophila* PLK4 dimerization mutant suggests that homodimerization is dispensable for its trans-phosphorylation (Ryniawec *et al*, 2023).

**Figure 1:**
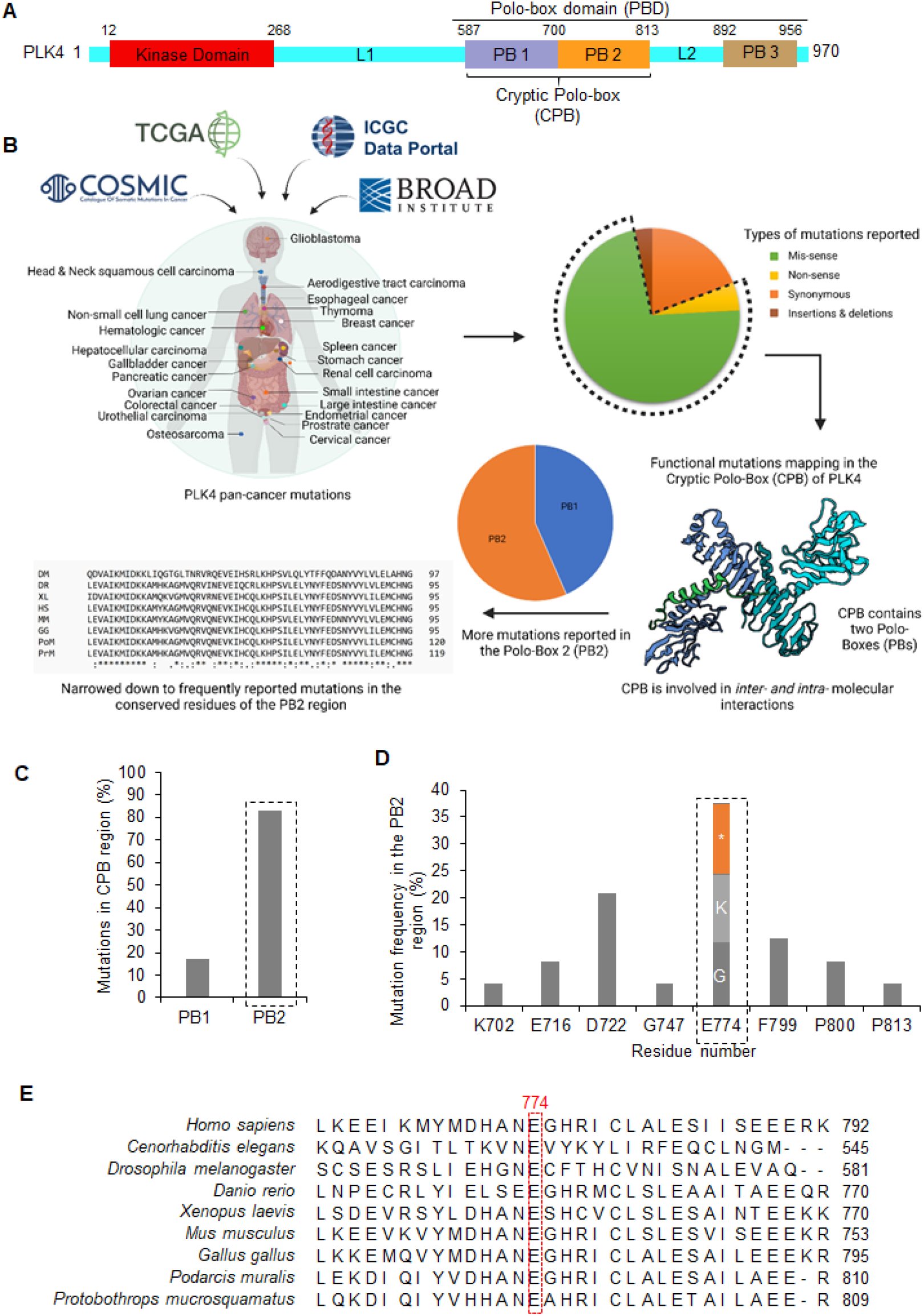
PLK4 E774 residue exhibits mutations in cancer patients. (**A**) Schematic representation of the conserved PLK4 domains. A number indicates amino acid residue. (**B**) Workflow for the identification of PLK4-associated functional mutations from pan-cancer datasets. Missense and nonsense mutations compiled from the indicated data portals were filtered based on their frequent occurrence at conserved residues in the cryptic polo-box (CPB) region of PLK4. (**C**) Bar graph indicating the frequency of mutation occurrence in polo-box 1 (PB1) and polo-box 2 (PB2) region of PLK4. (**D**) Bar graph showing the frequency of mutations at conserved residues of the PB2 region. The most frequently mutated glutamic acid (E) residue at the 774 position (E774) is either changed to Glycine (G), lysine (K), or truncated (*). (**E**) Multiple sequence alignment highlighting conservation of E774 residue across various indicated PLK4 homologs.

The 3D-structured illumination microscopy shows an early deposition of PLK4 dot at the site for the procentriole formation (Sonnen *et al*, 2012). In *C. elegans*, SPD-2, the human CEP192 orthologue (Delattre *et al*, 2006; Pelletier *et al*, 2006), and in *Drosophila*, Asterless (Asl), the human CEP152 orthologue (Dzhindzhev *et al*, 2010) are required for the recruitment of respective PLK4 orthologues (ZYG-1 in worms and SAK/PLK4 in flies) at the centrosomes. However, in vertebrates, both CEP192 and CEP152 are required for the PLK4 localization at the centrosome. The N-terminal region (1-217 residues) of CEP152 directly interacts with the CPB (589-887 residues) of PLK4 (Cizmecioglu *et al*, 2010; Hatch *et al*, 2010). Later, a competitive and mutually exclusive binding to CEP192 and CEP152, which works hierarchically, was found to be involved in PLK4 centriole recruitment (Kim *et al*, 2013). The super-resolution microscopy confirmed a unidirectional scaffold switching of PLK4 from CEP192 to CEP152 binding during the G_1_-phase of the cell cycle. Accordingly, an early ring of PLK4 colocalizes with CEP192, and the diameter of the ring increases when CEP152 appears at the centriole. The CPB binds with a 2:2 stoichiometry to the CEP192-58mer peptide and at a 2:1 stoichiometry to the CEP152-60mer peptide (Park *et al*, 2014). The findings suggest that CEP192 to CEP152 unidirectional scaffold could regulate PLK4 monomeric to dimeric transitioning. However, it is intriguing that in the unsynchronized cells, CEP152 knockdown increases PLK4 levels at centrosomes, whereas CEP192 inhibition decreases PLK4 centrosome levels (Cizmecioglu *et al*, 2010; Hatch *et al*, 2010; Sonnen *et al*, 2013). Therefore, the involvement of CEP192 to CEP152 unidirectional scaffold binding in regulating PLK4 centrosome recruitment requires better temporal understanding.

This work identified a cancer-linked PLK4 variant truncated at 774 residue position (E774*). This mutation affects PLK4 homodimerization and reveals a cross-dependency between PLK4 and CEP152 for localization at the S-phase centrosomes. Disturbance in the PLK4-CEP152 cross-dependency affects pericentrin levels at centrosomes, which results in unfocused spindles at the M-phase and reduces cell viability.

## RESULTS

### Identification of PLK4-associated cancerous mutations

We found a total of 1446 PLK4-associated amino acid mutations in the pan-cancer datasets collected from the publicly available databases, which include TCGA (https://www.cancer.gov/tcga), COSMIC (Tate *et al*, 2019), ICGC (Zhang *et al*, 2019), and Broad Institute (Ghandi *et al*, 2019). Among them, 78% (n=1127) were either missense or nonsense in nature, which were further narrowed down based on their occurrence in the cryptic polo-box (CPB) region (n=338) of PLK4. Further analysis revealed 29 functional mutations at conserved residues in the CPB region (**Fig 1B**). Among them, 83% were present in the polo-box 2 (PB2) within the CPB of PLK4 (**Fig 1C**). Interestingly, in 37.5% of total cases, there is a truncation at a conserved E774 residue in the PB2 (E774*), or the residue is substituted either by lysine or glycine (E774K or E774G) (**Fig 1D,E**). The E774 position was affected in patients suffering from uterine, kidney, endometrium, liver, thyroid, head and neck, skin, and aerodigestive tract cancers. It motivated us to investigate the influence of E774 residue changes on PLK4 functioning at centrosomes.

### PLK4 cancer-associated variant (E774*) affects its homodimerization

The CPB region is involved in PLK4 homodimerization (Leung *et al*, 2002; Slevin *et al*, 2012). We analyzed the X-ray crystal structure of human CPB from the protein data bank (PDB:4N7V) (Park *et al*, 2014), which suggests that the PLK4 cancerous mutation causing truncation of protein (E774*) disrupts the PB2 region involved in homodimerization (**Fig 2A**). The recombinant GST-tagged full-length PLK4 (GST-PLK4, 1-970 residues) was used to pull-down His-MBP labeled PLK4: PBD (571-970 residues), CPB (571-820 residues) or L2+PB3 (814-970 residues) (**Fig 2B**). The pull-down confirmed that the GST-PLK4 can dimerize with the CPB or the PBD, whereas the L2+PB3 region is not involved in dimerization (**Fig 2C**). Subsequently, we analyzed the effect of PLK4-associated cancerous mutants, i.e., nonsense (E774*) and missense (E774K) in the PB2 region on its homodimerization. The E774* mutation in PBD significantly reduces dimerization with GST-PLK4, whereas the E77K shows no effect on dimerization in the pull-down assay (**Fig 2D**). This suggests that in the case of E774* mutant PLK4 homodimerization is significantly affected, due to disruption of regions required for dimerization. Accordingly, we could show that deleting a stretch of 39 residues from 774-813 position in the PDB is suffient to loose PLK4 dimerization in the pull-down assay (**Fig 2E,F**).

**Figure 2:**
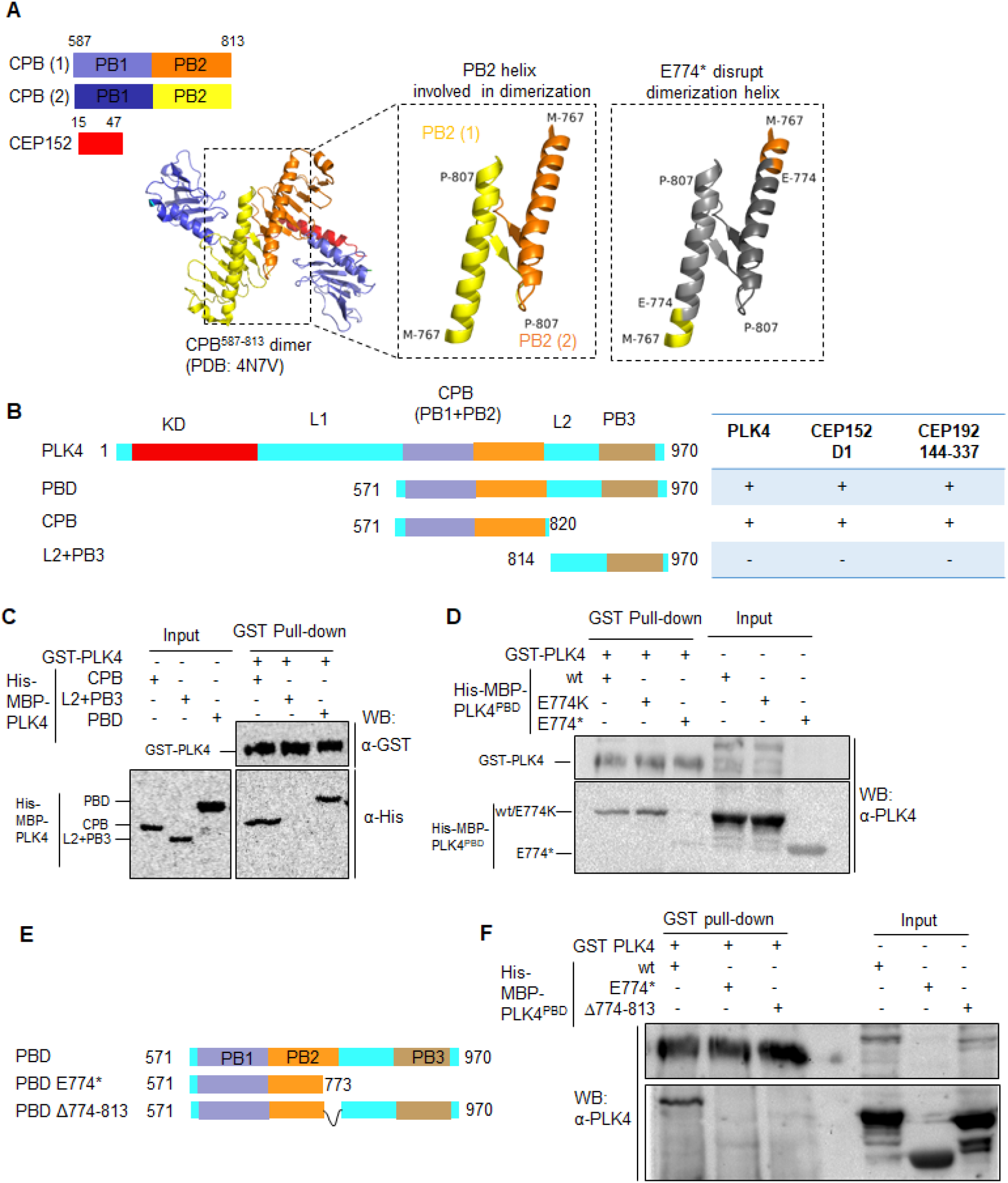
PLK4-associated cancerous mutation (E774*) disrupts homodimerization. **(A)** Structural analysis of X-ray diffraction structure from PDB:4N7V, representing PLK4 cryptic polo-box (CPB) dimer containing 767-807 residues. The magnified view highlights the involvement of the CPB region in dimer formation, which is disrupted by the E774* mutation. (B) Graphical summary of PLK4 domains and their interaction with GST-PLK4 (residues 1-970), GST-CEP152 domain 1(D1, residues 1-512) and GST-CEP192 (residues 144-337). (**C**) Western blot using antibodies against GST (α-GST) and His (α-His), representing interaction status between GST-PLK4 (residues 1-970) with the His-MBP PLK4: PBD (residues 571-970), CPB (residues 571-820) and L2+PB3 (residues 814-970) in the GST pull-down assay. (**D**) Western blot using antibodies against PLK4 (α-PLK4), representing the interaction status between GST-PLK4 with the His-MBP PLK4 PBD wild type (wt) and mutants (E774K and E774*) in the GST pull-down assay. (**E**) Schematic representation for PBD, truncation construct (PBD E774*), and deletion construct (PBD Δ774-813). (**F**) Western blot using an antibody against PLK4 (α-PLK4), representing interaction status between GST-PLK4 with the His-MBP PLK4 PBD wild type (wt) and mutants (E774* and Δ774-813) in the GST pull-down assay. Blots in C, D, and E are representations of at least two repeats.

### Cross-talk between CEP152 and PLK4 at the S-phase centrosomes

PLK4 is known to switch from CEP192 to CEP152 scaffold as the cell progresses from late G_1_ to S-phase, during the centrosome duplication (Kim *et al*, 2013). In human cells, CEP192-based regulations precede CEP152 (Sonnen *et al*, 2013). The analysis of CEP192 and CEP152 expression profiles indicates that both proteins remain confined to centrioles during early stages, but CEP192 spreads to expanding PCM as the cell progresses towards mitosis (Sonnen *et al*, 2013). Alternatively, the PLK4 levels at centrosomes are significantly less during interphase and reach their maximum levels at mitosis (Uchiumi *et al*, 1997).

We synchronized HeLa cells using a double thymidine block and analyzed cells at 0 hrs (Early S-phase) and 5 hrs post-release (Late S-phase) for the CEP192, CEP152, and PLK4 levels at centrosomes. Our analysis revealed that CEP192 protein levels are detectable at the early S-phase centrosomes but drop by the late S-phase. The PLK4 is detectable more at the late S-phase centrosomes when compared to the early S-phase. On the other hand, CEP152 levels are within detectable ranges at the early and late S-phase centrosomes (**Fig 3A-D**).

**Figure 3:**
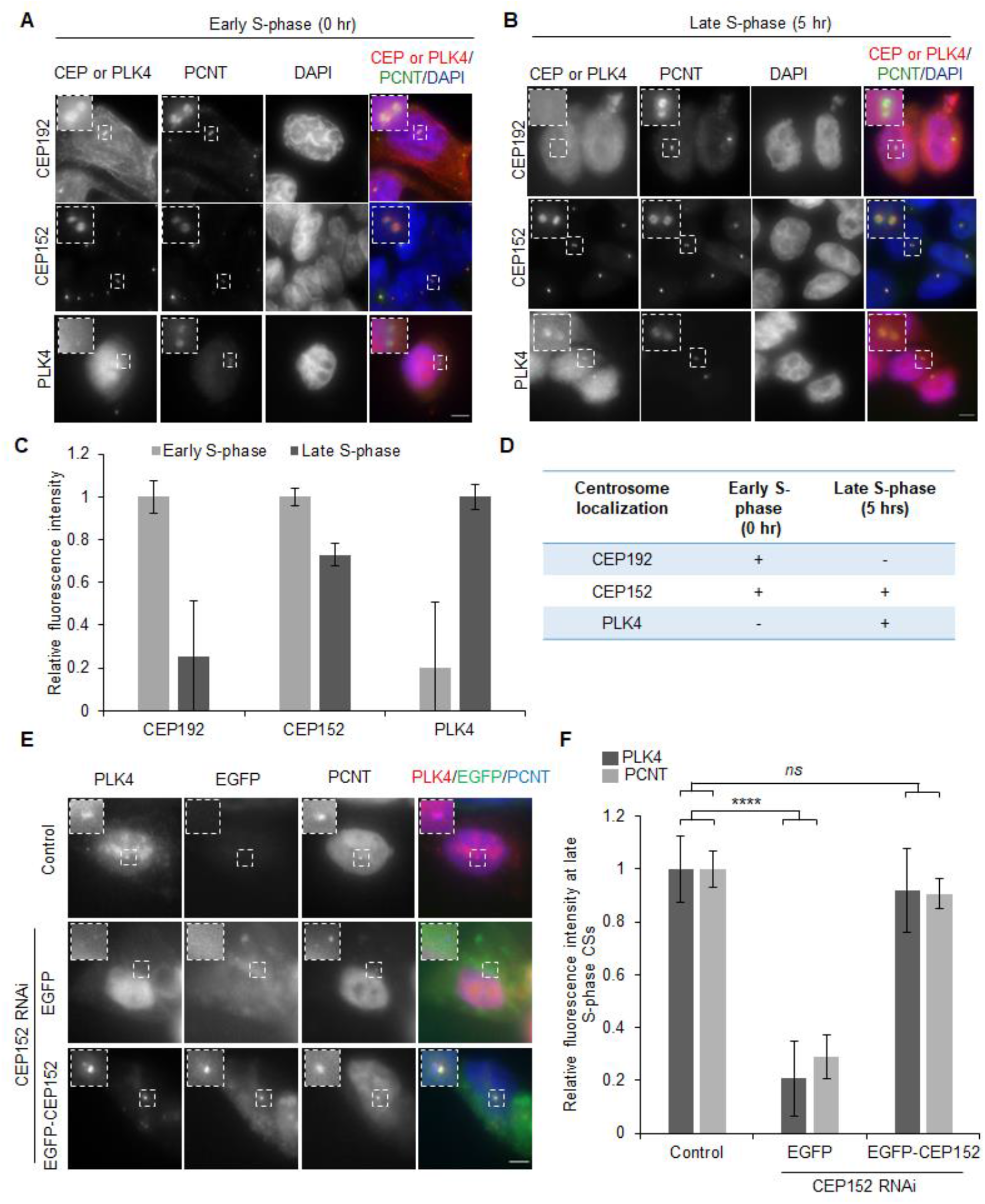
PLK4 localization at centrosomes is CEP152-dependent during the S-phase. (**A, B**) Representative immunofluorescence images of HeLa cells synchronized at (A) early S-phase and (B) late S-phase. Cells are immunostained with either of the CEP152,CEP192 or PLK4 (red), Pericentrin (PCNT) is used as a centrosome marker (green), and the nucleus is stained with DAPI (blue). Small inserts represent a maginified view of centrosomes in each panel. Scale bar: 5µm. (C) Bar graph showing relative fluorescence intensity at centrosomes ±SEM for CEP192 and CEP152 at late S-phase (dark grey) compared to early (light grey) S-phase cells. For PLK4, the relative fluorescence intensity at centrosomes±SEM at early (light grey) S-phase cells is shown compared to the late S-phase (dark grey). Results are from two independent experiments (n=40-100). (**D**) Table summarizing findings from A-C. (**E**) Representative immunofluorescence images of late S-phase HeLa cells immunostained for PLK4 (red) and PCNT (blue). EGFP (green) shows the signal from indicated expressed constructs. Inserts represent a magnified view of centrosomes for each panel. Scale bar: 5µm. (**F**) Bar graph showing relative fluorescence intensity at S-phase centrosomes for PLK4 (dark grey) and PCNT (light grey) compared to control. Results are from two independent experiments (n=30-50 cells). *ns*, not significant (*P>0.05*); \*\*\*\**P<0.0001* (two-tailed unpaired Student’s t-test)

CEP192 is known to be required for PLK4 localization at centrosomes (Kim *et al*, 2013; Sonnen *et al*, 2013), but the role of CEP152 is puzzling because its depletion increases PLK4 at centrosomes (Cizmecioglu *et al*, 2010; Hatch *et al*, 2010; Sonnen *et al*, 2013). To clarify the involvement of CEP152 in PLK4 recruitment, we focused on late S-phase centrosomes when both proteins are in detectable ranges. We depleted CEP152 using RNA interference (RNAi) (**Fig EV1A,B**), which significantly decreased (80%) endogenous PLK4 levels at the late S-phase centrosomes. Importantly, we were able to rescue PLK4 at centrosomes on expressing exogenous RNAi-resistant EGFP-CEP152, thus suggesting no off-target effects (**Fig 3E,F**). Our finding is in contrast to previous reports, where depletion of CEP152 in asynchronized U2OS cells resulted in increased PLK4 levels at the centrosome. Therefore, we also analyzed asynchronous U2OS cells to address this discrepancy and compared them to those synchronized at late S-phase. We observed increased endogenous PLK4 levels at centrosomes in asynchronized U2OS cells. However, cells synchronized at late S-phase corroborate our finding in HeLa cells (**Fig EV1C-F**). Therefore, we conclude that CEP152 is required for PLK4 centrosome localization at the late S-phase.

Next, we investigated the effect of PLK4 on the localization of CEP192 and CEP152 at the early and late S-phase centrosomes. PLK4 knockdown by RNA interference (RNAi) (**Fig EV1G,H**) did not affect CEP192 localization at centrosomes (**Fig EV2A,B**). However, CEP152 centrosome levels are significantly reduced at the early and late S-phase (**Fig EV2C,D**). This suggests feedback between CEP152 and PLK4 to maintain their centrosome levels during the S-phase

### Cross-dependency between PLK4 homodimerization and CEP152 for their centrosome localization

The CPB region of PLK4, involved in homodimerization, also interacts with the CEP192 (Kim *et al*, 2013) and CEP152 (Cizmecioglu *et al*, 2010; Hatch *et al*, 2010). We used GST-CEP192^144-337^ (residues 144-337) and GST-CEP152^D1^ (residues 1-512), which encompass residues required for interaction with the PLK4 CPB (Park *et al*, 2014). Accordingly, we could show that they were sufficient to pull-down the His-MBP tagged CPB and the PBD region, whereas L2+PB3 does not interact with the GST-CEP152^D1^ and GST-CEP192^144-337^ (**Figs 4A, EV3A**). The E774* homodimerization mutant could not interact with the CEP152^D1^ and CEP192^144-337^ in the GST pull-down assay (**Figs 4B, EV3B**). Previous work has shown the involvement of K685 and K711 residues in CEP152 interaction (Park *et al*, 2014). We also analyzed the X-ray crystal structure PDB:4N7V, which revealed the involvement of residues K685 in the PB1 and K711 in the PB2 region in direct interactions with CEP152 (**Fig 4C**). Accordingly, we showed that the PBD double mutant, K685/711A, can homodimerize (**Fig 4D**) but loses interaction with the GST-CEP152^D1^ compared to the wild-type PBD (**Fig 4E**). It suggests that the E774* mutant, which encompasses the CEP152 direct binding residues, possibly loses interaction with CEP152 due to its inability to homodimerize. The PBD (Δ774-813), which did not dimerize (**Fig 2E,F**), was also unable to interact with the GST-CEP152^D1^ (**Fig 4F,G**) or GST-CEP192^114-337^ (**Fig EV3B**), thus corroborating previous findings. Therefore, we conclude that PLK4 homodimerization contributes to CEP152 interaction.

**Figure 4:**
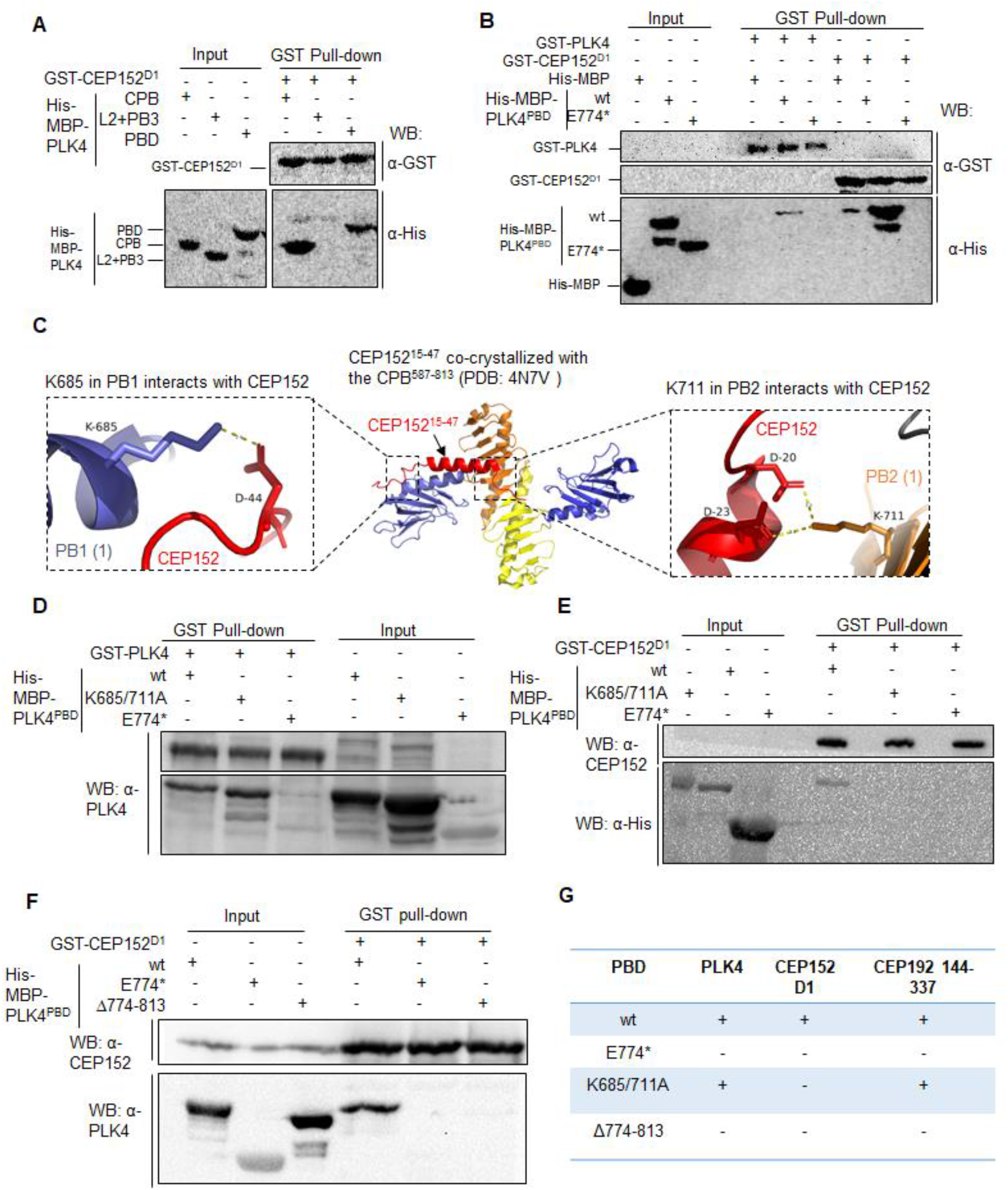
PLK4 homodimerization is involved in CEP152 interaction. (**A**) Western blot using antibodies against GST (α-GST) and His (α-His), representing interaction status between GST-CEP152 D1 (residues 1-512) and His-MBP tagged PLK4: PBD (residues 571-970), CPB (residues 571-820) and L2+PB3 (residues 814-970) using the GST pull-down assay. (**B**) Western blot representing interaction status of GST-PLK4 (residues 1-970) and GST CEP152 D1 with the His-MBP alone (control), His-MBP-PBD and His-MBP-PBD E774* mutant using the GST-pull down assay. (**C**) Structural analysis of PDB:4N7V crystal structure indicates the involvement of two residues, K685 in the PB1 and K711 in the PB2 region of PLK4 in interaction with CEP152. (**D, E**) Western blot representing interaction status of His-MBP tagged PBD, wild type (wt), double mutant (K685/711A), and E774* mutant with GST-PLK4 (D) and GST-52 D1 (E) using GST pull-down analysis. (**F**) Western blot representing interaction status of His-MBP tagged PBD: wild type (wt), E774*, and deletion (Δ774-813 residues) constructs with GST-152 D1 using GST pull-down analysis. (**G**) Table summarizing interaction between respective PBD constructs with PLK4 (for dimerization), CEP152 D1 (residues 1-512), and CEP192 (residues 144-337) proteins using GST-pull-down assays. Blots in A, B, D-F are representations of at least two repeats.

The above analysis indicates an effect of PLK4 homodimerization on CEP152 interaction. Next, we investigated the effect of PLK4 homodimerization on CEP152 centrosome levels at late S-phase. We observed about a 60% reduction in CEP152 centrosome levels on PLK4 depletion, which can be rescued by expressing RNAi-resistant exogenous EGFP-PLK4 (**Fig 5A,B**). Interestingly, homodimerization mutant EGFP-PLK4-E774* failed to rescue CEP152 levels at S-phase centrosomes (**Fig 5C,D**). The PLK4 E774* mutant expression is higher than the endogenous PLK4 (**Fig 5E**). Still, it shows reduced centrosome localization (**Fig 5F,G**), because it cannot interact with CEP152 and CEP192 (**Figs 4B, EV3B,C**).

**Figure 5:**
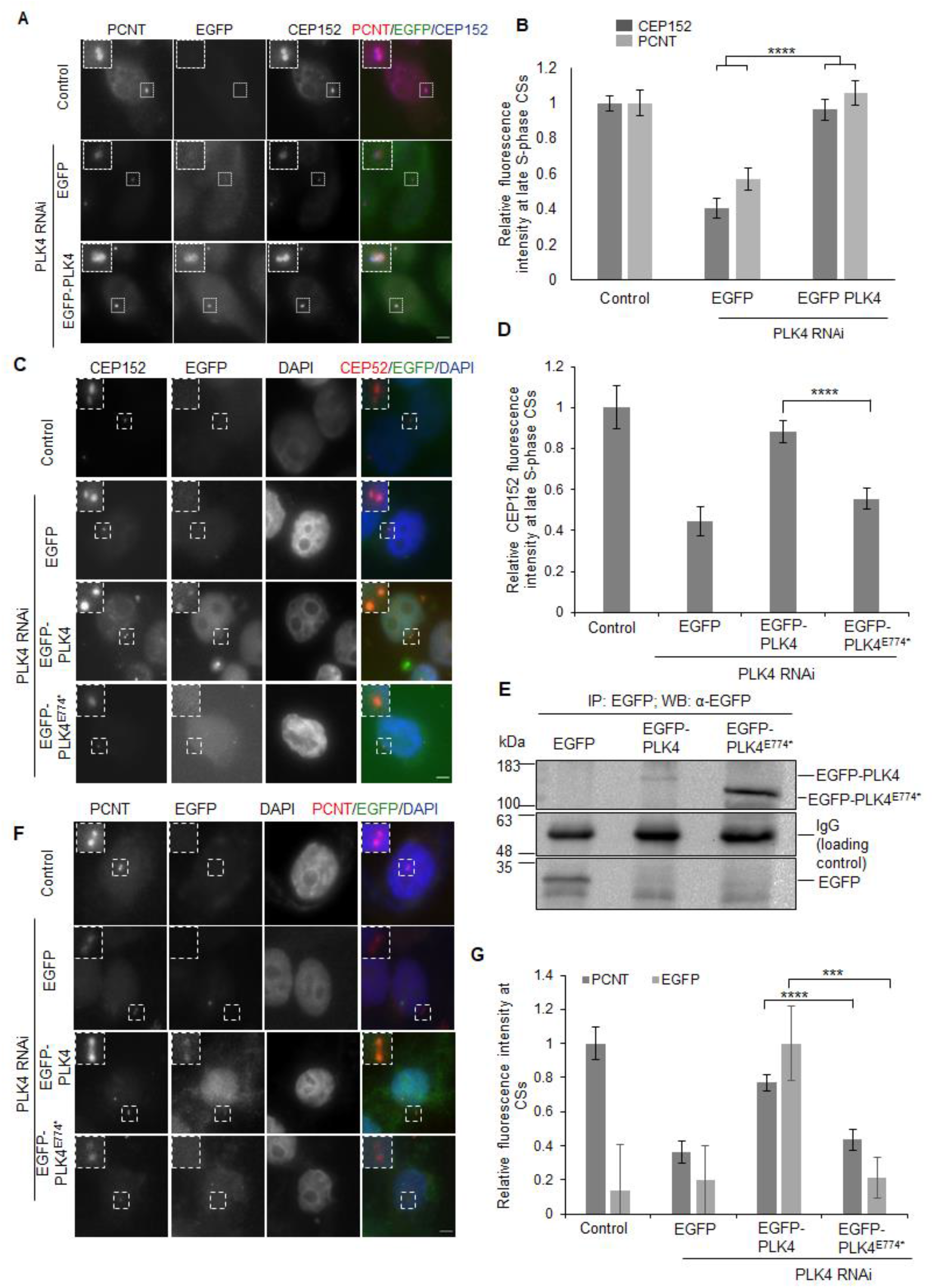
PLK4 homodimerization maintains centrosome levels of CEP152 and pericentrin. **(A)** Representative immunofluorescence images of HeLa cells immunostained with pericentrin (PCNT, red) and CEP152 (blue). EGFP (green) shows the signal from indicated expressed constructs. Inserts represent a magnified view of centrosomes in each panel. Scale bar: 5µm. **(B)** Bar graph showing relative CEP152 (dark grey) and PCNT (light grey) fluorescence intensity levels at late S-phase centrosomes±SEM compared to control. Results are from two independent experiments (n=40-70 cells). \*\*\*\**P<0.0001* (two-tailed unpaired Student’s t-test). (**C**) Representative immunofluorescence images of HeLa cells immunostained with CEP152 (red) and DAPI (blue). EGFP (green) shows the signal from indicated expressed constructs. Inserts represent a magnified view of centrosomes in each panel. Scale bar: 5µm. (**D**) Bar graph showing relative CEP152 fluorescence intensity levels at late S-phase centrosomes±SEM compared to control. Results are from two independent experiments (n=50-100 cells). \*\*\*\**P<0.0001* (two-tailed unpaired Student’s t-test). (**E**) Western blot showing immunoprecipitated (IP) proteins, EGFP, EGFP-PLK4, and EGFP-PLK4 E774* from the transfected HeLa cell using the antibody against EGFP. IgG bands serve as a loading control. The blot is representative of two repeats. (**F**) Representative immunofluorescence images of HeLa cells immunostained with PCNT (red) and DAPI (blue). EGFP (green) shows the signal from indicated expressed constructs. Inserts represent a magnified view of centrosomes in each panel. Scale bar: 5µm. (**G**) Bar graph showing relative PCNT (dark grey) fluorescence intensity levels at late S-phase centrosomes±SEM compared to control and relative EGFP (light grey) fluorescence intensity levels at late S-phase centrosomes±SEM compared to the condition in which endogenous PLK4 is depleted by RNA interference (RNAi) and complemented with transfected RNAi-resistant EGFP-PLK4. Results are from two independent experiments (n=50-100 cells). \*\*\**P<0.001* and \*\*\*\**P<0.0001* (two-tailed unpaired Student’s t-test).

Previously reported K685 residue is involved in interaction with CEP192 (Park *et al*, 2014), but mutating it to alanine in the double mutant (K685/711A) did not affect interaction with GST-CEP192^144-337^ in the pull-down assay (**Fig EV3C**). The EGFP-PBD double mutant (K685/711A) mutant had similar localization as wild-type PBD at the S-phase centrosomes when the endogenous PLK4 was depleted by siRNA (**Fig EV3D,E**), which could be due its ability to homodimerize and maintain CEP192 interaction. Notably, the K685/711A PBD mutant, despite losing direct CEP152 interaction (**Fig 4E**), can rescue CEP152 levels at centrosomes without endogenous PLK4 (**Fig EV3D,E**). The data strongly suggest that PLK4 dimerization is required for its centrosome localization and maintaining CEP152 levels of centrosomes at S-phase.

### CEP152-PLK4 feedback maintains spindle organization and cell viability

Previous work using CRISPR/Cas9-mediated CEP152 depletion showed that it affects pericentrin levels in interphase cells (Watanabe *et al*, 2019). We also observed a correlation between CEP152 and pericentrin levels at centrosomes (**Fig EV1A,B**). Accordingly, depletion of PLK4 or rescue with the homodimerization mutant E774*, which significantly decreases CEP152 levels at centrosomes, resulted in a significant reduction of pericentrin levels at centrosomes (**Fig 5 A,B,F,G**). Pericentrin is essential for spindle organization (Purohit *et al*, 1999; Chen *et al*, 2014). Subsequently, we observed a significant increase in the population of cells with unfocused spindles at the M-phase of the cell cycle when the endogenous PLK4 was knocked down by siRNA. The effect can be rescued on the expression of EGFP-PLK4. However, the EGFP-PLK4 E774* homodimerization mutant could not rescue the phenotype (**Fig 6A,B**). Since problems in spindle organizations could affect cell proliferation and viability (Levine & Holland, 2018).

**Figure 6:**
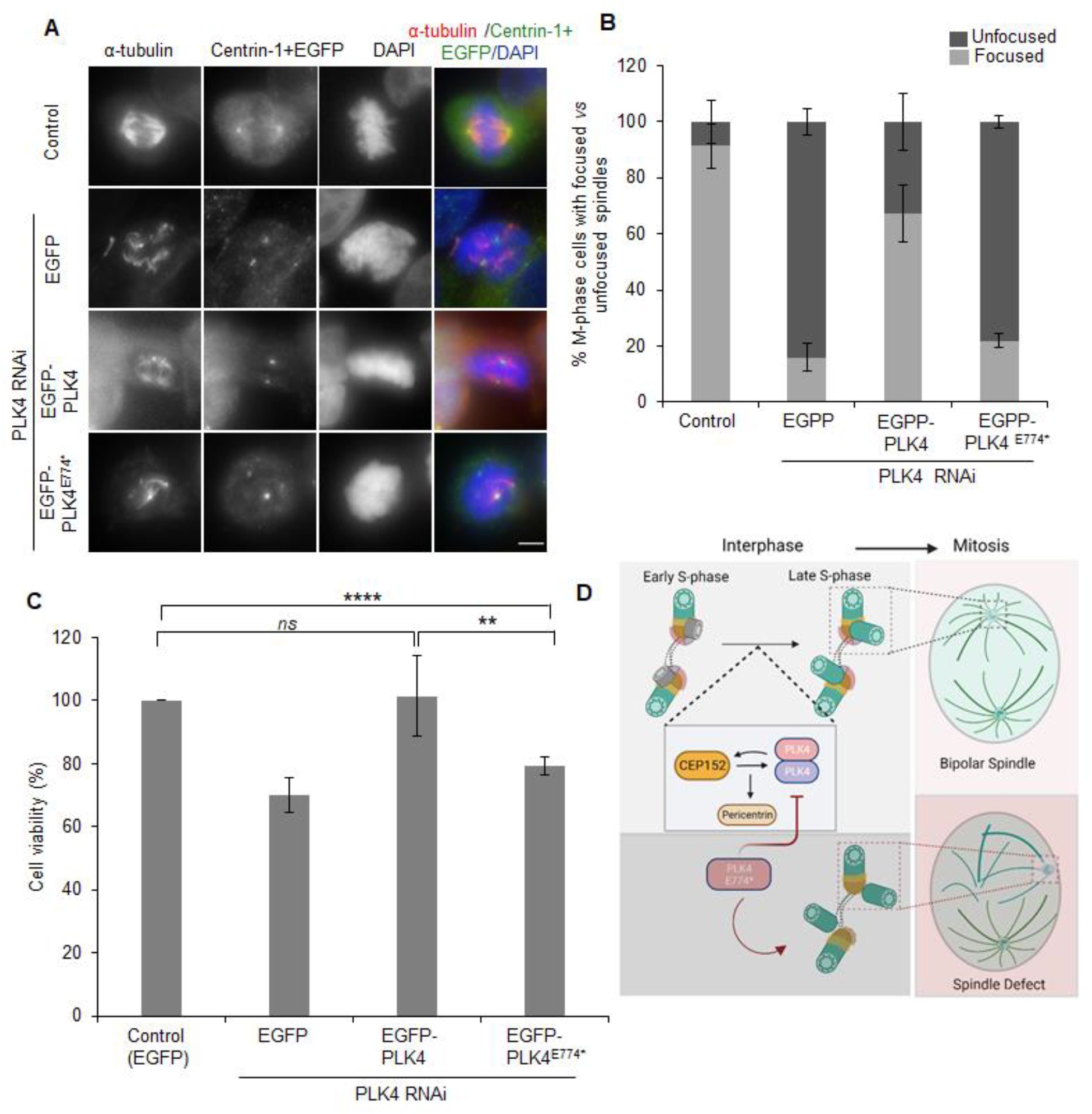
PLK4 homodimerization mutant affects spindle organization at M-phase and cell viability. (**A**) Representative images of HeLa cells immunostained with α-tubulin (red), centrin-1+EGFP signal (green), and DAPI (blue). Scale bar: 5µm. (**B**) Bar graph representing average percentage (%) of mitotic cells with focused (light grey) and unfocussed (dark grey) spindles±SD under indicated conditions. Results are from two independent experiments (n=30-60 cells). (**C**) Bar graph representing average percentage (%) of cell viability±SD for respective conditions using the MTT assay. Results are from triplicates for each state repeated in four independent experiments. **(D**) The proposed model depicts cross-dependency between CEP152 and PLK4 homodimerization for their centrosome localization during the S-phase progression. The CEP152-PLK4 cross-talk regulates pericentrin levels at the centrosome, which generate focused spindles at the M-phase. The PLK4-associated cancerous variant E774* causes loss of PLK4 homodimerization and disturbs CEP152-PLK4 dependency. It consequently reduces pericentrin levels and results in unfocused spindles.

We then performed the MTT-based cell viability assay, which confirmed a significant decrease in the viability of cells on PLK4 depletion or rescue with the homodimerization mutant E774* (**Fig 6C**). Thus, we conclude that PLK4 homodimerization is required for proper centrosome and spindle organization in human cells, which also has implications for their viability.

## DISCUSSION

The PLK4 Cryptic Polo-Box (CPB) region containing two polo-boxes (PB1 and PB2) in tandem is required for homodimerization, and the same region is also involved in interaction with the upstream centrosome recruiters, i.e., CEP192 and CEP152. PLK4 homodimerization activates trans-phosphorylation of regulatory elements in the linker-1 (L1) region, which are recognized by the SLIMB-SCF^βTrCP^-dependent ubiquitination and subjected to protein degradation. In *Drosophila* PLK4, dimerization relieves the autoinhibitory folding of L1 region on the kinase domain and thus assists in trans-activation (Klebba *et al*, 2015). However, a mutation in fly PLK4, which inhibits protein dimerization, does not affect PLK4 activation or centriole localization (Ryniawec *et al*, 2023). The mechanisms can differ in complex organisms because human PLK4 transitions from CEP192 to CEP152 scaffold as the cell progresses from G_1_ to S-phase (Park *et al*, 2014). CEP192 is required for PLK4 recruitment as inhibition of CEP192 reduces PLK4 levels at centrosomes (Kim *et al*, 2013; Sonnen *et al*, 2013). However, the involvement of CEP152 in PLK4 recruitment is puzzling, as previous literature reported that CEP152 depletion causes an increase in PLK4 levels at centrosomes (Cizmecioglu *et al*, 2010; Hatch *et al*, 2010; Sonnen *et al*, 2013). In this work, we carefully analyzed the CEP192-CEP152-PLK4 network in cells synchronized at early and late S-phase. Our analysis revealed that CEP192 centrosome levels remain unaffected by PLK4 depletion. However, there is a cross-dependency of PLK4 and CEP152 for their centrosome localization at the S-phase. We identified PLK4-associated frequently occurring cancerous variant (E774*) mapping in the PB2 region of protein from the pan-cancer datasets. The E774* disrupts CPB dimerization, which could be verified using the *in vitro* pull-down assay. Our work further highlights the involvement of a 39 amino acid stretch (residues 774-813) in the PB2 region of CPB sufficient for the dimerization of PLK4.

Interestingly, this also affects direct binding with the CEP152, despite maintaining residues required for CEP152 interaction (K685 and K711), thus pointing towards the need of homodimerization for CEP152 interaction. Although K685 residue in the PB1 of PLK4 is required for CEP192 interaction in the immunoprecipitation assay (Park *et al*, 2014), the double mutant (K685/711A) showed no impact on the direct CEP192 binding in the pull-down assay. Later analysis showed proper centrosome localization of K685/711K comparable to the wild-type PLK4 protein. It suggests that posttranslational modifications inside the cell might influence PLK4 and CEP192 interaction.

Importantly, we could show that the cancerous variant of PLK4 (E774*) cannot maintain CEP152 levels at S-phase centrosomes. The loss of centrosome CEP152 in PLK4 RNAi or E774* rescue cells correlates with a decrease in pericentrin levels at centrosomes. Since pericentrin is required for generating focused spindles in mitotic cells, we could show an increase in the population of cells with unfocused spindles under conditions resulting in pericentrin reduction (**Fig 6D**). This finding suggests that human PLK4 homodimerization is required for CEP152-mediated centriole organization and downstream events involved in spindle organization. Disruption of PLK4 homodimerization is associated with cancer. Therefore, understanding the requirement of PLK4 dimerization for regulating functional aspects of centrosome organization is relevant for developing potential therapeutic strategies.

## MATERIAL & METHODS

### In silico analysis

PLK4 amino acid variations were compiled from pan-cancer datasets available at TCGA (https://www.cancer.gov/tcga), COSMIC (Tate *et al*, 2019), ICGC (Zhang *et al*, 2019), and Broad Institute (Ghandi *et al*, 2019). Missense and nonsense mutations in the PLK4 CPB region were considered for analysis. Multiple sequence alignment of PLK4 homologous sequences from different classes of the animal kingdom (Mammalia, Insecta, Amphibia, Rodent, Actinoptergii, Aves, Reptilia, and Nematoda) was performed using Clustal Omega (Goujon *et al*, 2010; Sievers *et al*, 2011) tool.

The X-ray diffracted structure of human PLK4 cryptic polo-box and CEP152: 4N7V (Park *et al*, 2014) from the Protein Data Bank was analyzed using PyMOL (Schrödinger, LLC, 2015) protein visualization tool. Distance between amino acid residues is measured using the distance measurement wizard tool in PyMOL.

### Plasmids and cloning

PLK4 cDNA (gift from Prof. Andrea Musacchio) coding for full-length and different domains were PCR amplified. The PLK4 coding regions were cloned in pETDuet-1 plasmid for 6xHis-MBP tag (gift from Prof. Andrea Musacchio), pGEX-6P-1 for GST tag (gift from Prof. Andrea Musacchio) and pcDNA5/FRT/TO plasmid for EGFP tag (gift from Prof. Andrea Musacchio) at the N-terminus of the protein. The amplified products were ligated between the *BamH*1 (Takara, #1010A) and the *Xho*1 (Takara, #094A) restriction sites in the same reading frame. For cloning CEP152 D1 (residues 1-512), the corresponding coding region was PCR amplified (CEP152 cDNA gifted from Prof. Pierre Gönczy) and ligated in pGEX-6P-1 plasmid downstream of GST using the *BamH1* and *Xho1* restriction sites. CEP192 144-337 region was synthesized and cloned in pGEX-6P-1 plasmid for the GST tag at N-terminus (Biomatik, Canada). Site-directed mutations were generated using specific primers for amplification with Phusion High-fidelity DNA polymerase (New England Biolabs, #M0530S), followed by template digestion with *DpnI* restriction enzyme (Takara, #1235A). The amplified product was transformed in *Escherichia coli* NEB5α cells (NEB#C2987) to obtain colonies. All plasmids were validated by the Sanger sequencing method.

### Protein expression and GST pull-down assay

*Escherichia coli* BL21 (gift from Prof. Andrea Musacchio) strain cells were used for bacterial protein expression. The bacterial cultures were grown in an orbital shaker until OD_600_ of 0.5-0.7 at 37°C. Proteins were induced using 0.5–1 mM IPTG for 16-18 hrs at 18°C.

The bacterial cells expressing GST-tagged proteins were resuspended in GST lysis buffer (50mM Hepes pH 7.5, 150mM NaCl, 1% Triton-X, 1mM DTT, 10% Glycerol, 1mM PMSF) and sonicated. The supernatant collected after centrifugation was incubated with pre-equilibrated glutathione superflow resin (Takara, #635607) for 2-3 hrs at 4°C on continuous shaking. Beads were collected at 300 g for 2 min and washed thrice.

The His-tagged proteins were resuspended in the lysis buffer (50mM Tris-HCl pH 8.0, 100mM NaCl, 5mM Imidazole, 10% Glycerol, 1mM PMSF) and sonicated. The supernatant collected after centrifugation was incubated with the glutathione superflow resin-bound GST-tagged protein for 1 hr at 4°C with continuous shaking. The beads were washed 4-5 times with GST lysis buffer, and the protein complex was eluted in the Laemmli sample buffer. The protein samples were boiled and resolved on 8% SDS-PAGE.

### Western blotting

For western blotting, resolved protein bands from SDS-PAGE were transferred to the nitrocellulose membrane. The membrane was blocked using 5% skimmed milk in 1X PBST (1X PBS with 0.05% Tween-20). The blot was incubated with a primary antibody prepared in a blocking buffer. It was washed thrice with 1XPBST and incubated with an HRP-conjugated secondary antibody. After final washing, the blot was developed using the Clarity western ECL substrate (Biorad, #1705061). The SDS-PAGE gel and blot images were taken using the Gel documentation system (Biotron Omega Lum G).

### Cell culture

HeLa Kyoto and U2OS cells (gifts from Prof. Andrea Musacchio), were cultured in High glucose DMEM medium (Himedia, #AL111) supplemented with 10% fetal bovine serum (Himedia, #RM10432), 2mM L-Glutamine (Sigma, #G7513) and 2mM penicillin and streptomycin antibiotics (Sigma, #P4333) in a 5% CO_2_ containing humidified incubator (Eppendorf Galaxy 170S) at 37°C.

### Cell transfection, cell synchronization, and siRNA

According to the manufacturer’s protocol, plasmids and siRNA were transfected in mammalian cells at 70% confluency using Lipofectamine 3000 (Invitrogen, #L3000001). PLK4 siRNA and CEP152 siRNA oligonucleotides (Merck) were synthesized with 3’UU overhangs for the following sense strand sequences 5-’CUCCUUUCAGACAUAUAAG-3’ (Ohta *et al*, 2018) and 5’-GGAGGACCAUGUCAUUAGACU-3’, respectively. Cells were synchronized to the G_1_/S boundary with a double thymidine block (Chen & Deng, 2018). Briefly, 2mM thymidine (Sigma #T9250) was added to the cells for 18 hrs, followed by release in fresh media for 9 hrs. Next round of thymidine block using 2mM thymidine was provided for 15 hrs. For early S-phase, cells were processed just after the second thymidine block. For the late S-phase, cells are released for 5 hrs in fresh media before analyzing, and for the M-phase, cells were released for 12 hrs.

### Primary and Secondary antibodies

Primary antibodies used in western blotting (WB) and immunofluorescence (IF) were anti-6x His mouse monoclonal (Invitrogen, #MA1-21315, 1:5000 in WB), anti-GST mouse monoclonal (Invitrogen, #MA4-004, 1:5000 in WB), anti-PLK4 rabbit polyclonal (Abcam, #ab137398, 1:2000 in WB, 1:50 in IF), anti-CEP152 rabbit polyclonal (Abcam, #ab-183911, 1:2500 in WB, 1:200 in IF), anti-GFP rabbit polyclonal (Sigma, #G1544, 1:7000 in WB), anti-CEP192 rabbit polyclonal (Abcam, #ab229705, 1:20 in IF), anti-pericentrin mouse monoclonal (Abcam, #ab28144, 1:100 in IF), anti-α tubulin mouse monoclonal (Sigma, #T9026, 1:100 in IF) and anti-centrin-1 rabbit polyclonal (Abcam, #ab156858, 1:200 in IF).

Secondary antibodies used in western blotting were anti-mouse IgG HRP-linked antibody (Cell Signaling Technology, #7076S, 1:10,000) and anti-rabbit IgG HRP-linked antibody (Cell Signalling Technology, #7074S, 1:10,000). Secondary antibodies used in immunofluorescence were anti-rabbit IgG–Rhodamine goat polyclonal (Merck, #SAB3700846, 1:500), anti-mouse IgG– Rhodamine rabbit polyclonal (Merck, #SAB3701020, 1:500), anti-rabbit IgG–FITC antibody sheep polyclonal (Merck, #F7512, 1:750), anti-mouse IgG–FITC antibody goat polyclonal (Merck, #F5387, 1:750), anti-rabbit IgG (H+L) cross-adsorbed Alexa Fluor 405 goat polyclonal (Invitrogen, #A-31556, 1:250) and anti-mouse IgG (H+L) cross-adsorbed Alexa Fluor 405 goat polyclonal (Invitrogen, #A-31553, 1:250).

### Immunoprecipitation

Cells were resuspended in the RIPA lysis buffer (10mM Hepes pH 7.5, 50mM NaCl, 0.5% Sodium deoxycholate, 0.1% SDS, 0.5% NP40) along with 1mM PMSF and 100 ul/ml protease inhibitor cocktail. The cell suspension was passed through a small syringe a few times and incubated on ice for 30 min. The supernatant was collected after centrifugation and incubated with a specific primary antibody overnight at 4°C. The pre-equilibrated protein G-sepharose 4B conjugate beads (Invitrogen, #101241) were then added to the sample and incubated for 4 hrs at 4°C. Beads were collected and washed thrice using RIPA buffer. The bound protein is eluted in the Laemmli sample buffer and analyzed by western blotting.

### Immunofluorescence

Hela Kyoto or U2OS cells were grown on coverslips coated with Poly-D lysine (Merck, #P7280). Coverslips containing cells were gently washed twice with 1X PBS to remove excess media. Cells are permeabilized using 0.1% Triton X-100 (Himedia, #TC286) in 1X PBS, followed by a fixing step using 4% paraformaldehyde (Sigma, #158127). After washing them with 1XPBS, cells were blocked in 1% BSA (Himedia, #MB083) solution in 1x PBS with 0.1% Tween-20 (Himedia, #MB067). Primary antibody was added over coverslips and incubated overnight. Cells were washed thrice with 1X PBS and incubated with a secondary antibody conjugated to a fluorescent probe. The excess antibody was removed by washing with 1x PBS, and the nucleus was stained using DAPI (0.5ug/ml) prepared in 1x PBS. After final washing, the coverslip was mounted on a glass slide using mounting solution (0.62 g DABCO, 2 g Mowiol, 6.25 ml glycerol, 25 ml 0.2 M Tris-Cl pH 8.5). The slides were visualized using the fluorescence microscope’s 100X oil immersion, 1.35 NA objective lens (Olympus IX83). Z-stacks with 0.25 um distance are captured. Signal intensity measurements and localizations were analyzed using ImageJ (Schindelin *et al*, 2012) software.

### Fluorescence Intensity measurement at centrosomes

Pericentrin (PCNT) was used as a marker to locate centrosomes, except in Fig 5,C,D and Fig EV E,D, where the CEP152 signal was used. Using ImageJ, a circular region of interest (ROI) was drawn at PCNT, and the same ROI was copied near the centrosome, which served as background fluorescence. The same ROIs were duplicated in the other channels to measure their centrosome fluorescence intensity and background. Absolute fluorescence intensity at the centrosome was calculated by subtracting the background signal. The average fluorescence intensity at centrosomes relative to control±Standard Error of the Mean (SEM) was plotted using Microsoft Excel.

### MTT assay

About 5000 cells per well were seeded in a 96-well plate (Himedia, #TPP96) and allowed to settle. Cells were transfected using PLK4 siRNA. The next day, cells were transfected with the EGFP-expressing plasmids. On the third day, 0.5 mg/ml MTT (3-(4,5-dimethylthiazol-2-yl)-2,5-diphenyl tetrazolium bromide; Invitrogen,#M6494) solution prepared in serum free-media was added to the cells and incubated for 4 hrs at 37°C in a humidified CO_2_ incubator. Formazan crystals were dissolved using 200ul/well of acidified isopropanol under continuous shaking. The absorbance at 570nm was measured using a microplate reader (Biotek synergy H1). Bar graph representing the percentage of cell viability was plotted in Microsoft Excel.

### Statistical analysis and Softwares

Experiments were performed at least three times, and more than 50 cells were used for fluorescence intensity measurement analysis. Fluorescence signal intensity measurements were performed using ImageJ (Schindelin *et al*, 2012). Bar graphs were plotted in Microsoft Excel. A two-tailed unpaired Student’s t-test was used for statistical analysis and indicated on top of bar graphs using a solid line between the two compared conditions. Figure panels were arranged on Microsoft PowerPoint.

## DATA AVAILABILITY

This study includes no data deposited in external repositories

## ACKNOWLEDGEMENTS

The HeLa Kyoto, U2OS, *Escherichia coli* BL21, pET Duet1, pGEX 6P1, pCDNA5 FRT/TO, and PLK4 cDNA were kind gifts from the laboratory of Prof. Andrea Musacchio, Max Planck Institute of Molecular Physiology, Dortmund, Germany. CEP152 cDNA was a kind gift from the laboratory of Prof. Pierre Gönczy, Swiss Federal Institute of Technology Lausanne (EPFL), Lausanne, Switzerland. We thank the Centre for Research & Development of Scientific Instruments at IIT Jodhpur for the microscopy facility.

## FUNDING

H.K. is supported by a fellowship (09/1125(0017)/2020-EMR-I) from the Council of Scientific & Industrial Research, Government of India. The work is supported by the grants received from the Department of Biotechnology (BT/12/IYBA/2019/02), and the Board of Research in Nuclear Sciences (55/14/02/2021-BRNS/10206).

## DISCLOSURE & COMPETING INTERESTS STATEMENT

The authors declare no competing or financial interests.

## AUTHOR CONTRIBUTIONS

Conceptualization: H.K., P.S.; Methodology: H.K.; Validation: H.K., P.S.; Formal analysis: H.K., P.S.; Investigation: H.K.; Data curation: H.K., P.S.; Writing original draft: H.K.; Writing-review & editing: P.S.; Supervision: P.S.; Project administration: P.S.; Funding acquisition: P.S.

---

**Figure EV1:**
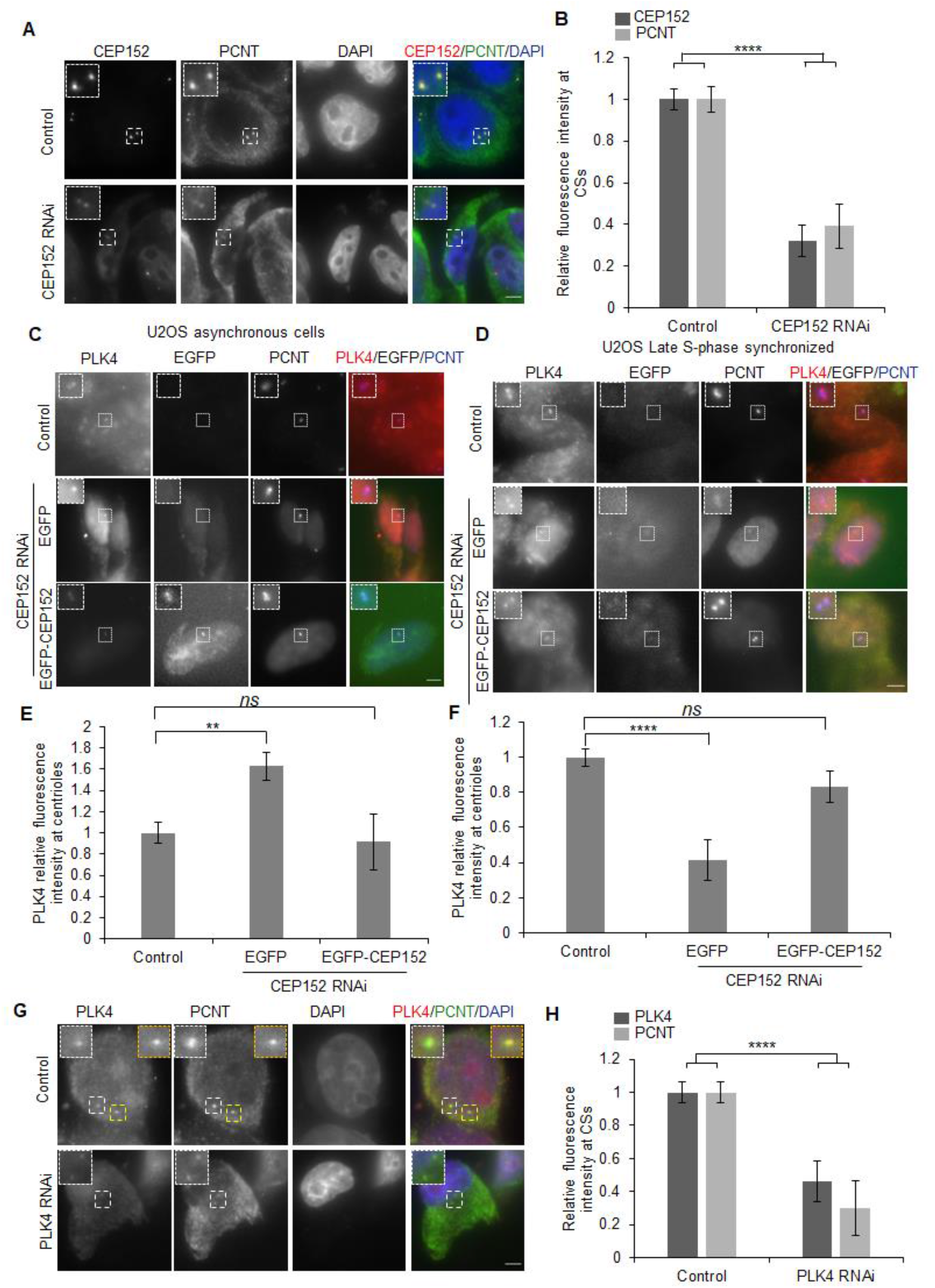
(**A**) Representative immunofluorescence images of HeLa cells immunostained for CEP152 (red), pericentrin (PCNT, green), and DAPI (blue) for control and CEP152 knocked-down using small interfering RNA (RNAi). Inserts in each panel represent a maginified view of centrosomes. Scale bar: 5µm. (**B**) Bar graph showing relative fluorescence levels of CEP152 (dark grey) and PCNT (light grey) ±SEM compared to control cells (n=50-100 cells). \*\*\*\**P<0.0001* (two-tailed unpaired Student’s t-test). (**C,D**) Representative immunofluorescence images of U2OS cells immunostained for PLK4 (red) and PCNT (blue), which expresses EGFP or RNAi-resistant EGFP-CEP152 when the endogenous CEP152 is depleted in asynchronous (C) and late S-phase (D) synchronized cells. Inserts represent a magnified view of centrosomes. Scale bar: 5µm. (**E, F**) Bar graph showing relative levels of PLK4 at centrosomes±SEM for asynchronous (E) and PCNT late S-phase (F) synchronized cells from the experiment (C, D). Results are from two independent experiments (n=30-100 cells). *ns* (not significant, \*\**P<0.01* and \*\*\*\**P<0.0001* (two-tailed unpaired Student’s t-test). (**G**) Representative immunofluorescence images of HeLa cells immunostained for PLK4 (red), PCNT (green), and DAPI (blue) for control and PLK4 knocked-down using small interfering RNA (PLK4 RNAi). Inserts represent a magnified view of centrosomes. Scale bar: 5µm. (**H**) Bar graph showing relative levels of PLK4 (dark grey) and PCNT(light grey) at centrosomes±SEM (n = 40-70 cells). \*\*\*\**P<0.0001* (two-tailed unpaired Student’s t-test).

**Figure EV2:**
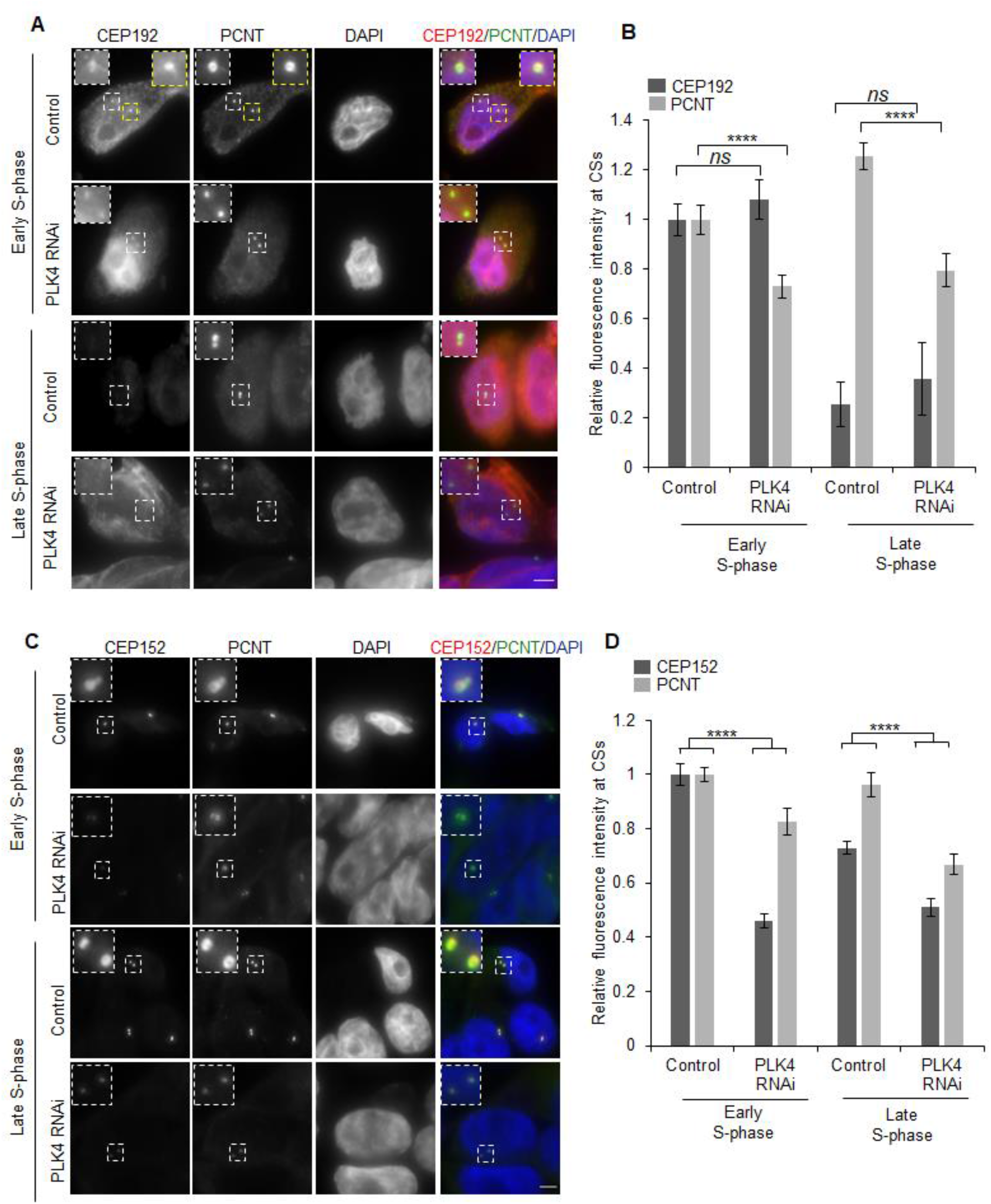
(**A, C**) Representative immunofluorescence images of HeLa cells immunostained for (A) CEP192 or (C) CEP152 (red), PCNT (green), and DAPI (blue) in control and PLK4 RNAi at the early and the late S-phase. Inserts represent a magnified view of centrosomes in each panel. Scale bar: 5µm. (**B, D**) Bar graph showing relative levels of (B) CEP192 or (D) CEP152 (dark grey) and PCNT (light grey) at the centrosomes±SEM (n=40-100 cells). *ns* (not significant, \*\*\*\**P<0.0001* (two-tailed unpaired Student’s t-test).

**Figure EV3:**
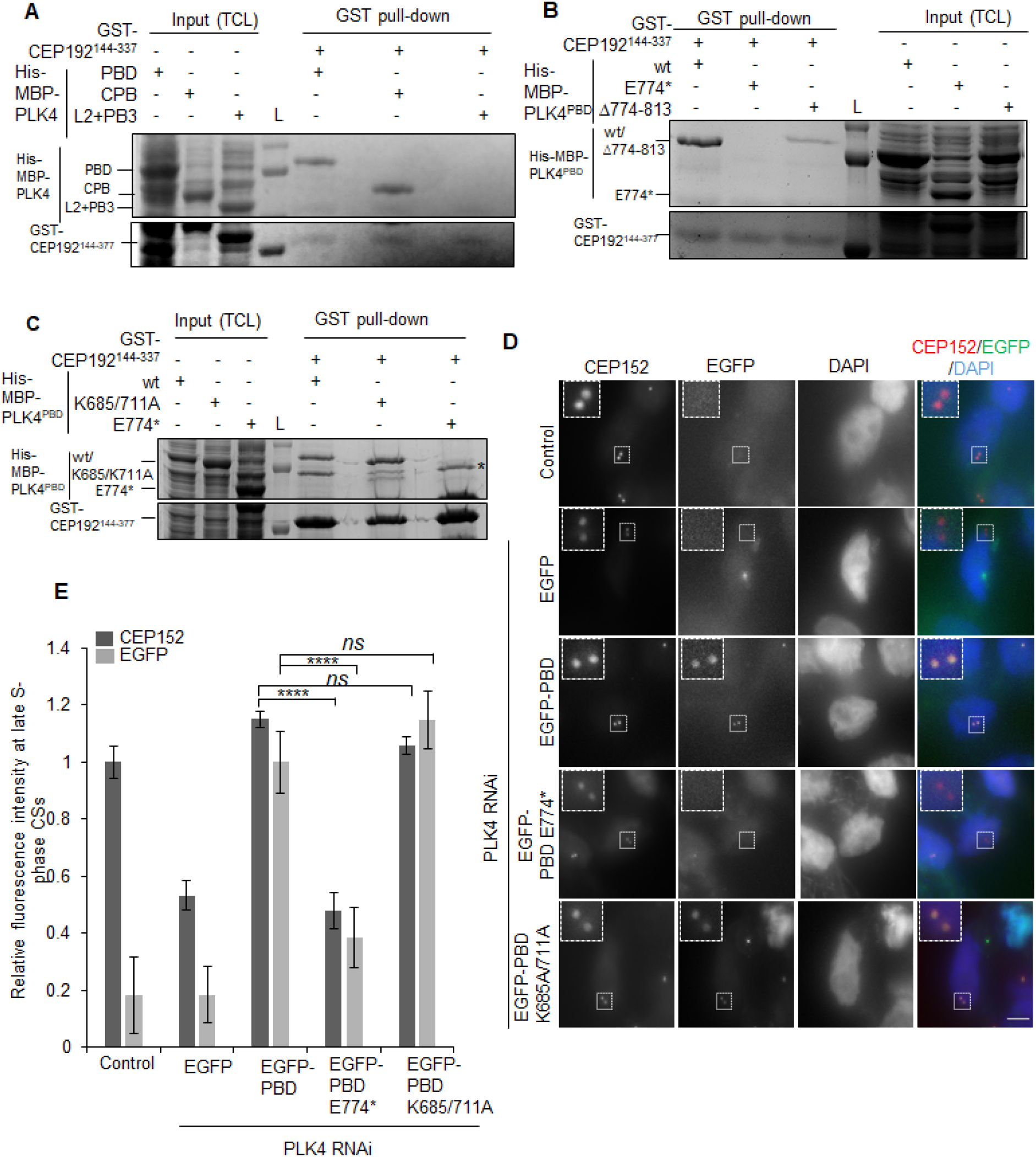
(**A-C**) Coomassie-stained SDS-PAGE gels showing indicated inputs from total cell lysates (TCL), and GST pull-down assays using GST tagged CEP192 having residues 144-377 and showing interaction status with the His-MBP PLK4 related domains and mutants. The gels are representative of at least three repeats. *non–specific band (**D**) Representative immunofluorescence images of HeLa cells immunostained for CEP152 (red) and DAPI (blue). The cells express EGFP or EGFP-PLK4: PBD, PBD E774*, and PBD K685/711A mutant in PLK4 RNAi background. Inserts represent a magnified view of centrosomes. Scale bar: 5µm. (**E**) Bar graph showing relative levels of CEP152 and EGFP at the late S-phase centrosomes±SEM (n=50-100 cells). *ns* (not significant, \*\*\*\**P<0.0001* (two-tailed unpaired Student’s t-test).

## Notes

### Competing Interest Statement

The authors have declared no competing interest.

### Summary of Updates

This version of the manuscript has been revised to update the expanded view figures

## REFERENCES

Barr FA, Silljé HHW & Nigg EA (2004) Polo-like kinases and the orchestration of cell division. Nat Rev Mol Cell Biol 5: 429–441

Bettencourt-Dias M, Rodrigues-Martins A, Carpenter L, Riparbelli M, Lehmann L, Gatt MK, Carmo N, Balloux F, Callaini G & Glover DM (2005) SAK/PLK4 is required for centriole duplication and flagella development. Curr Biol 15: 2199–2207

Chen C-T, Hehnly H, Yu Q, Farkas D, Zheng G, Redick SD, Hung H-F, Samtani R, Jurczyk A, Akbarian S, et al (2014) A unique set of centrosome proteins requires pericentrin for spindle-pole localization and spindle orientation. Curr Biol 24: 2327–2334

Chen G & Deng X (2018) Cell synchronization by double thymidine block. Bio-protoc 8

Cizmecioglu O, Arnold M, Bahtz R, Settele F, Ehret L, Haselmann-Weiß U, Antony C & Hoffmann I (2010) Cep152 acts as a scaffold for recruitment of Plk4 and CPAP to the centrosome. J Cell Biol 191: 731–739

Cunha-Ferreira I, Bento I, Pimenta-Marques A, Jana SC, Lince-Faria M, Duarte P, Borrego-Pinto J, Gilberto S, Amado T, Brito D, et al (2013) Regulation of autophosphorylation controls PLK4 self-destruction and centriole number. Curr Biol 23: 2245–2254

Delattre M, Canard C & Gönczy P (2006) Sequential protein recruitment in C. elegans centriole formation. Curr Biol 16: 1844–1849

Dzhindzhev NS, Yu QD, Weiskopf K, Tzolovsky G, Cunha-Ferreira I, Riparbelli M, Rodrigues-Martins A, Bettencourt-Dias M, Callaini G & Glover DM (2010) Asterless is a scaffold for the onset of centriole assembly. Nature 467: 714–718

Fu J, Hagan IM & Glover DM (2015) The centrosome and Its duplication cycle. Cold Spring Harb Perspect Biol 7: a015800

Fujita H, Yoshino Y & Chiba N (2015) Regulation of the centrosome cycle. Mol Cell Oncol 3: e1075643

Ganem NJ, Godinho SA & Pellman D (2009) A mechanism linking extra centrosomes to chromosomal instability. Nature 460: 278–282

Ghandi M, Huang FW, Jané-Valbuena J, Kryukov GV, Lo CC, McDonald ER, Barretina J, Gelfand ET, Bielski CM, Li H, et al (2019) Next-generation characterization of the cancer cell line encyclopedia. Nature 569: 503–508

Godinho SA & Pellman D (2014) Causes and consequences of centrosome abnormalities in cancer. Philos Trans R Soc Lond B Biol Sci 369: 20130467

Goujon M, McWilliam H, Li W, Valentin F, Squizzato S, Paern J & Lopez R (2010) A new bioinformatics analysis tools framework at EMBL–EBI. Nucleic Acids Res 38: W695–W699

Habedanck R, Stierhof Y-D, Wilkinson CJ & Nigg EA (2005) The Polo kinase Plk4 functions in centriole duplication. Nat Cell Biol 7: 1140–1146

Hatch EM, Kulukian A, Holland AJ, Cleveland DW & Stearns T (2010) Cep152 interacts with Plk4 and is required for centriole duplication. J Cell Biol 191: 721–729

Jaiswal S, Kasera H, Jain S, Khandelwal S & Singh P (2021) Centrosome: A Microtubule Nucleating Cellular Machinery. J Indian Inst Sci 101: 5–18

Kim T-S, Park J-E, Shukla A, Choi S, Murugan RN, Lee JH, Ahn M, Rhee K, Bang JK, Kim BY, et al (2013) Hierarchical recruitment of Plk4 and regulation of centriole biogenesis by two centrosomal scaffolds, Cep192 and Cep152. Proc Natl Acad Sci U S A 110: E4849–4857

Klebba JE, Buster DW, McLamarrah TA, Rusan NM & Rogers GC (2015) Autoinhibition and relief mechanism for Polo-like kinase 4. Proc Natl Acad Sci 112: E657–E666

Klebba JE, Buster DW, Nguyen AL, Swatkoski S, Gucek M, Rusan NM & Rogers GC (2013) Polo-like kinase 4 autodestructs by generating its Slimb-binding phosphodegron. Curr Biol 23: 2255–2261

Kleylein-Sohn J, Westendorf J, Le Clech M, Habedanck R, Stierhof Y-D & Nigg EA (2007) Plk4-induced centriole biogenesis in human cells. Dev Cell 13: 190–202

Leung GC, Hudson JW, Kozarova A, Davidson A, Dennis JW & Sicheri F (2002) The Sak polo-box comprises a structural domain sufficient for mitotic subcellular localization. Nat Struct Biol 9: 719–724

Levine MS & Holland AJ (2018) The impact of mitotic errors on cell proliferation and tumorigenesis. Genes Dev 32: 620–638

Liu L, Zhang CZ, Cai M, Fu J, Chen GG & Yun J (2012) Downregulation of polo-like kinase 4 in hepatocellular carcinoma associates with poor prognosis. PLoS One 7: e41293

Marina M & Saavedra HI (2014) Nek2 and Plk4: prognostic markers, drivers of breast tumorigenesis and drug resistance. Front Biosci (Landmark Ed) 19: 352–365

Ohta M, Watanabe K, Ashikawa T, Nozaki Y, Yoshiba S, Kimura A & Kitagawa D (2018) Bimodal Binding of STIL to Plk4 Controls Proper Centriole Copy Number. Cell Rep 23: 3160–3169.e4

Park S-Y, Park J-E, Kim T-S, Kim JH, Kwak M-J, Ku B, Tian L, Murugan RN, Ahn M, Komiya S, et al (2014) Molecular basis for unidirectional scaffold switching of human Plk4 in centriole biogenesis. Nat Struct Mol Biol 21: 696–703

Pelletier L, O’Toole E, Schwager A, Hyman AA & Müller-Reichert T (2006) Centriole assembly in Caenorhabditis elegans. Nature 444: 619–623

Purohit A, Tynan SH, Vallee R & Doxsey SJ (1999) Direct Interaction of Pericentrin with Cytoplasmic Dynein Light Intermediate Chain Contributes to Mitotic Spindle Organization. J Cell Biol 147: 481–492

Ryniawec JM, Buster DW, Slevin LK, Boese CJ, Amoiroglou A, Dean SM, Slep KC & Rogers GC (2023) Polo-like kinase 4 homodimerization and condensate formation regulate its own protein levels but are not required for centriole assembly. Mol Biol Cell: mbcE22120572

Schindelin J, Arganda-Carreras I, Frise E, Kaynig V, Longair M, Pietzsch T, Preibisch S, Rueden C, Saalfeld S, Schmid B, et al (2012) Fiji: an open-source platform for biological-image analysis. Nat Methods 9: 676–682

Schrödinger, LLC (2015) The PyMOL Molecular Graphics System, Version 1.8.

Shimanovskaya E & Dong G (2014) Expression, purification and preliminary crystallographic analysis of the cryptic polo-box domain of Caenorhabditis elegans ZYG-1. Acta Crystallogr F Struct Biol Commun 70: 1346–1350

Shimanovskaya E, Viscardi V, Lesigang J, Lettman MM, Qiao R, Svergun DI, Round A, Oegema K & Dong G (2014) Structure of the C. elegans ZYG-1 cryptic polo box suggests a conserved mechanism for centriolar docking of Plk4 kinases. Structure 22: 1090–1104

Shinmura K, Kurabe N, Goto M, Yamada H, Natsume H, Konno H & Sugimura H (2014) PLK4 overexpression and its effect on centrosome regulation and chromosome stability in human gastric cancer. Mol Biol Rep 41: 6635–6644

Sievers F, Wilm A, Dineen D, Gibson TJ, Karplus K, Li W, Lopez R, McWilliam H, Remmert M, Söding J, et al (2011) Fast, scalable generation of high-quality protein multiple sequence alignments using Clustal Omega. Mol Syst Biol 7: 539

Slevin LK, Nye J, Pinkerton DC, Buster DW, Rogers GC & Slep KC (2012) The structure of the Plk4 cryptic polo box reveals two tandem polo boxes required for centriole duplication. Struct 20: 1905–1917

Sonnen KF, Gabryjonczyk A-M, Anselm E, Stierhof Y-D & Nigg EA (2013) Human Cep192 and Cep152 cooperate in Plk4 recruitment and centriole duplication. J Cell Sci 126: 3223–3233

Sonnen KF, Schermelleh L, Leonhardt H & Nigg EA (2012) 3D-structured illumination microscopy provides novel insight into architecture of human centrosomes. Biol Open 1: 965–976

Tate JG, Bamford S, Jubb HC, Sondka Z, Beare DM, Bindal N, Boutselakis H, Cole CG, Creatore C, Dawson E, et al (2019) COSMIC: the Catalogue Of Somatic Mutations In Cancer. Nucleic Acids Res 47: D941–D947

Uchiumi T, Longo DL & Ferris DK (1997) Cell cycle regulation of the human Polo-like Kinase (PLK) promoter. J Biol Chem 272: 9166–9174

Wang J, Zuo J, Wang M, Ma X, Gao K, Bai X, Wang N, Xie W & Liu H (2019) Polo–like kinase 4 promotes tumorigenesis and induces resistance to radiotherapy in glioblastoma. Oncol Rep 41: 2159–2167

Watanabe K, Takao D, Ito KK, Takahashi M & Kitagawa D (2019) The Cep57-pericentrin module organizes PCM expansion and centriole engagement. Nat Commun 10: 931

Zhang J, Bajari R, Andric D, Gerthoffert F, Lepsa A, Nahal-Bose H, Stein LD & Ferretti V (2019) The International Cancer Genome Consortium Data Portal. Nat Biotechnol 37: 367–369

